# Astrocyte-derived thrombospondin induces cortical synaptogenesis in a sex-specific manner

**DOI:** 10.1101/2021.01.04.425242

**Authors:** Anna Mazur, Ean H. Bills, Brandon J. Henderson, W. Christopher Risher

## Abstract

The regulation of synaptic connectivity in the brain is vital to proper functioning and development of the central nervous system (CNS). Formation of neural networks in the CNS has been shown to be heavily influenced by astrocytes, which secrete factors, including thrombospondin (TSP) family proteins, that promote synaptogenesis. However, whether this process is different between males and females has not been thoroughly investigated. In this study, we found that cortical neurons purified from newborn male rats showed a significantly more robust synaptogenic response compared to female-derived cells when exposed to factors secreted from astrocytes. This difference was driven largely by the neuronal response to TSP2, which increased synapses in male neurons while showing no effect on female neurons. Blockade of endogenous 17β-estradiol production with letrozole normalized the TSP response between male and female cells, indicating a level of regulation by estrogen signaling. Our results suggest that TSP-induced synaptogenesis is critical for the development of male but not female cortical synapses, contributing to sex differences in astrocyte-mediated synaptic connectivity.

## Introduction

Neurons form complex arrangements throughout the central nervous system (CNS) that form the basis of our ability to think, move, learn, and remember. Synapses, the basic functional units of the CNS, represent the coming together of an axon from a presynaptic cell and a dendrite from a postsynaptic one. Precise timing and location of synapse formation (i.e. synaptogenesis) is critical for the function of the developing brain, and it has become increasingly apparent that astrocytes are powerful regulators of this process (Barres, 2008). The last twenty-plus years have seen an explosion of discovery of astrocytic mechanisms that influence synaptogenesis, both positively and negatively, via both contact-mediated signaling as well as secreted factors (Risher and Eroglu, 2020). However, despite this greatly increased awareness, a significant caveat of most of these studies was the lack of consideration for potential sex differences in astrocyte-mediated synaptic development.

The brains of males and females show significant structural, chemical, and functional differences (Pfaff and Joe◻ls, 2017). This phenomenon holds true over the course of “normal” development as well as in cases of aberrant CNS function, such as that seen in neurodevelopmental and neurodegenerative disorders (Hanamsagar and Bilbo, 2016). The release of steroids from the gonads beginning with puberty certainly plays a major role in defining sex differences in the still-maturing brain (Genazzani et al., 2007; Ho et al., 2020), but relatively little is known about how pre-pubertal cellular processes may influence neural circuits in a distinct, sex-dependent manner. Many previous investigations into astrocyte-induced synaptogenesis were performed on neurons isolated from mixed-sex litters of rats and mice, often utilizing astrocyte-conditioned media (ACM) or astrocyte culture inserts which were similarly generated from mixed-sex sources (Allen et al., 2012; Blanco-Suarez et al., 2018; Christopherson et al., 2005; Eroglu et al., 2009; Farhy-Tselnicker et al., 2017; Kucukdereli et al., 2011). However, in our recent study elucidating the cellular mechanism underlying cortical synaptogenesis via the astrocyte-secreted matricellular protein, thrombospondin (TSP), we switched to using only male mice and rats for our neuronal purifications (Risher et al., 2018). The reason for this switch lies in our (previously unpublished) observations that purified TSP only increased excitatory synapses between cultured cortical neurons ~50% of the time when using mixed-sex litters. Conversely, neurons isolated from only males responded positively to TSP with a nearly 100% success rate, necessitating further investigation into this novel sex-dependent mechanism.

In this study, we have identified differences in astrocyte-mediated synapse formation between male and female-derived neurons. Neuronal cultures isolated from the cortices of male rats experienced a significantly greater fold-increase in excitatory synapses when exposed to astrocyte-secreted factors than did cultures purified from females. Driving this sex difference was the strongly synaptogenic response of male-derived neurons to TSP, which was not observed in female cultures. Intriguingly, our data suggest that the divergence may be, at least in part, due to transient differences in *de novo* neuronal estrogen synthesis during the peak synaptogenic window for astrocytes and TSP.

## Methods

### Animals

All experiments were conducted in accordance with Marshall University’s Institutional Animal Care and Use Committee (IACUC) guidelines (W.C.R. protocol 698). Adult Sprague-Dawley rat mothers and their mixed sex newborn litters were obtained from Hilltop Lab Animals. Following sex determination (based on visual examination of the anogenital region), pups were separated into male and female prior to dissection and cell purification.

### Cortical neuron purification and treatment

Cortical neurons were purified separately from male and female postnatal day 1 (P1) Sprague-Dawley rats by sequential immunopanning as previously described (Risher et al., 2018). Briefly, following rapid decapitation, brains were removed and cortices were dissected. Following 45 minutes of digestion in papain (~7.5 units/mL; Worthington) supplemented with DNase (Worthington) dissolved in Dulbecco’s PBS (DPBS; Gibco) at 34°C, sequential low/high concentrations of ovomucoid inhibitor (Worthington) was used to gradually halt papain activity. The cell solution was then passaged through a 20 μm Nitex mesh filter (Sefar) prior to a series of negative immunopanning steps. Petri dishes coated with Bandeiraea Simplicifolia Lectin I (Vector Laboratories), followed by AffiniPure goat-anti mouse IgG+IgM (H+L) (Jackson ImmunoResearch Laboratories) and AffiniPure goat-anti rat IgG+IgM (H+L) (Jackson ImmunoResearch Laboratories) antibodies were used to remove non-neuronal cells and general debris. To further purify neurons (>95%) using positive immunopanning, the cell solution was passaged onto petri dishes coated with mouse antibody against neural cell adhesion molecule L1 (ASCS4; Developmental Studies Hybridoma Bank, Univ. Iowa). Following final washes, pelleting and resuspension in serum-free neuronal growth medium (GM) [Neurobasal (Gibco), B27 supplement (Gibco), 2 mM GlutaMax (Gibco), 100 U/mL Pen/Strep (Gibco), 1 mM NaPyruvate (Gibco), 50 ng/mL BDNF (Peprotech), 20 ng/mL CNTF (Peprotech), 4.2 μg/mL forskolin (Sigma), and 10 μM AraC (Sigma)], neurons were plated at a density of 60K/well on poly-D-lysine (PDL; Sigma) and laminin (R&D Systems)-coated coverslips in a 24-well plate as previously described (Risher et al., 2018). After 2 days *in vitro* (DIV 2) at 37°C/5% CO_2_, half of the GM in each well was replaced with fresh GM of the same composition with the exception of: Neurobasal Plus (Gibco) instead of Neurobasal and GlutaMax, B27 Plus (Gibco) instead of B27, and no AraC; this media was then used for feedings every 2-3 days for the duration of the experiment prior to electrophysiology or fixation for synaptic immunocytochemistry (ICC) on DIV 13. Recombinant TSP2 protein was purified from CHO cells expressing mouse TSP2 via affinity chromatography with HiTRAP heparin HP (GE Healthcare) following protocols described by Oganesian et al. (2008). Standard TSP2 treatment of neurons was performed on DIVs 7 and 10 at a dose of 500 ng/mL.

### Cortical astrocyte isolation, culture, and astrocyte-conditioned media (ACM) harvesting

Cortical astrocytes were purified separately from male and female P1 Sprague-Dawley rats following a similar protocol to neurons, as described above. After the Nitex mesh filtering step, the cells were pelleted and resuspended in astrocyte growth media [AGM; DMEM + GlutaMax (Gibco), 10% heat-inactivated FBS (Sigma), 10 μM hydrocortisone (Sigma), 100 U/mL Pen/Strep, 5 μg/mL Insulin (Sigma), 1 mM NaPyruvate, 5 μg/mL N-Acetyl-L-cysteine (NAC; Sigma)]. After counting, 15-20 million cells were plated on PDL-coated 75mm^2^ flasks and incubated at 37°C/5% CO_2_. On DIV3, AGM was removed and replaced with DPBS. In order to isolate the adherent monolayer of astrocytes, flasks were shaken vigorously by hand 3-6x for 15 seconds each. DPBS was then replaced with fresh AGM. AraC was added to the AGM from DIV 5 to 7 to minimize astrocyte proliferation. On DIV 7, astrocytes were passaged into either transwell inserts (Corning) at a density of 125K (for direct culture with neurons) or at 3 million per 100 mm tissue culture dish (for the generation of ACM).

For the generation and collection of astrocyte-conditioned media (ACM), at 24 hrs post-passaging into 100 mm dishes, astrocytes were washed 3x with DPBS and conditioned at 37°C/5% CO_2_ without disturbing for 4 days in minimal medium [Neurobasal (phenol red-free), 2 mM L-Glutamine (Gibco), 1 mM NaPyruvate, 100 U/mL Pen/Strep, 5 μg/mL NAC, 40 ng/mL T3 (Sigma), and Sato supplement (Winzeler and Wang, 2013)]. After conditioning, media was collected from dishes and centrifuged to pellet cell debris prior to concentrating in 5,000 M.W.C.O. Vivaspin tubes (Sartorius). ACM was then aliquoted and flash frozen in liquid nitrogen until use. Standard ACM treatment of neurons was performed on DIVs 7 and 10 at a dose of 75 μg/mL prior to fixation for synaptic ICC on DIV 13.

### Synaptic immunocytochemistry (ICC)

Synapse quantification of cortical neurons followed previously established protocols (Kucukdereli et al., 2011; Risher et al., 2018; Risher et al., 2014). Briefly, DIV 13 neurons in 24 well plates were fixed with warm 4% paraformaldehyde (PFA) (Electron Microscopy Sciences) in PBS for 7 minutes. Next, PFA was removed and cells were washed 3x with PBS then blocked in a buffer containing 0.2% Triton X-100 (Roche) in 50% normal goat serum (NGS; Jackson ImmunoResearch)/50% antibody buffer (PBS containing 1% BSA, 0.04% NaN3, 0.2% Triton X-100) at room temperature for 30 minutes. Following another 3x PBS wash, cells were treated with 10% NGS/90% antibody buffer containing primary antibodies against Bassoon (1:500; mouse; Enzo/Assay Designs) and Homer1 (1:500; rabbit; Synaptic Systems), at 4°C overnight in the dark. The next morning, another 3x PBS wash was performed, followed by a 2 hour room temperature incubation in 10% NGS/90% antibody buffer containing the following fluorescently-conjugated secondary antibodies: Goat anti-mouse Alexa 488 (1:500; Invitrogen) and goat anti-rabbit Alexa 594 (1:500; Invitrogen). After final round of 3x PBS washes, coverslips were transferred to glass slides with Vectashield mounting medium containing DAPI (Vector Laboratories), sealed with nail polish, and imaged on a Leica DM5500B microscope with a 63x/1.4 NA objective at 1920×1440 resolution.

### Whole-cell patch clamp electrophysiology

Glass coverslips with neuronal cultures were detached from imaging dishes and placed in the recording chamber of an upright microscope (Axio Examiner, Zeiss) while being continually perfused by extracellular solution (ECS, in mM: 140 NaCl, 5 KCl, 2 CaCl_2_, 1 MgCl_2_, 10 HEPES, and 10 glucose). Electrophysiological signals were recorded from visually identified neurons with a Sutter IPA and SutterPatch software. Patch pipettes were filled with solution containing (in mM): 135 K gluconate, 5 KCl, 5 EGTA, 0.5 CaCl_2_, 10 HEPES, 2 Mg-ATP, and 0.1 GTP (pH was adjusted to 7.2 with Tris-base, and osmolarity was adjusted to 280-300 mOsm with sucrose). The resistance of patch pipettes was 4-8 MΩ. Junction potential was nulled just before forming a gigaseal. Series resistance was monitored without compensation throughout the experiment. Data were discarded if the series resistance (10-25 MΩ) changed by more than 20% during recordings. All recordings were done at room temperature. After achieving a gigaseal, gentle suction was used to achieve whole cell configuration. The neuron was voltage clamped at −70 mV to record miniature excitatory postsynaptic currents (mEPSCs) in the presence of 1.0 μM tetrodotoxin. Data were sampled at 10 kHz and filtered at 2 KHz.

### Enzyme-linked immunosorbent assay (ELISA) for estradiol

Cortical neurons isolated separately from male and female P1 Sprague-Dawley rats were plated at 60K/well into 48-well plates coated with PDL and laminin as described previously. Astrocytes were isolated from the same litter as described above, then plated at 600K/well into 6-well plates on DIV 7. Starting on DIV 1 (neurons) or DIV 5 (astrocytes), medium was collected daily, two wells pooled and flash-frozen per time point, until DIV 12. Concentration of estradiol in collected medium was then determined with Parameter Estradiol assay (R&D Systems) according to instructions provided by the manufacturer. Briefly, medium was thawed on ice, centrifuged to precipitate cellular debris, and incubated in ELISA plate wells coated with primary antibody against estradiol, in the presence of fixed amount of HRP-conjugated estradiol. Next, the absorbance was read at 450nm and corrected by reading at 570nm. Data were analyzed using the four-parameter logistic (4-PL) curve-fit, per manufacturer recommendations, generated with Prism software (GraphPad).

### Image Analysis and Statistics

Synapse quantification was performed using a custom Puncta Analyzer plugin for ImageJ (NIH) by a trained analyst who was blinded as to condition/treatment. The plugin allows for rapid counting of pre- (i.e. Bassoon), post- (i.e. Homer1), and co-localized synaptic puncta, determined by user-defined thresholds for each individual channel. This approach provides an accurate estimation of synapse number based on the precise localization of pre- and postsynaptic proteins, which are typically non-overlapping when imaged with fluorescence-based ICC except when directly opposed at synapses (Ippolito and Eroglu, 2010; Risher et al., 2018; Risher et al., 2014).

Briefly, the plugin separates raw images into “green” (488/Bassoon) and “red” (594/Homer1) channels, subtracts background (rolling ball radius=50), and then prompts the analyst to define a threshold with the goal of isolating “true” puncta while keeping noise to a minimum (which is facilitated by setting a minimum pixel size value of 4 in the user-selectable settings). The plugin then displays a results window with numbers of pre-, post- and co-localized synaptic puncta that can be copied to a Microsoft Excel spreadsheet. In order to calculate “% of GM” values, co-localized puncta values for the same sex growth media-only (GM) condition were averaged, then values across all treatments/conditions were converted to % of the calculated GM average.

Statistical analyses were performed in Microsoft Excel and Graphpad Prism. All data are presented as mean ± SEM. Statistical analyses were conducted using either unpaired t-tests, two-way ANOVAs or linear mixed-effects model [REML] with Holm-Sidak’s multiple comparisons *post hoc* test. Significance was shown as *p<0.05, **p<0.01, ****p<0.0001. Values that did not attain significance were either not noted or shown as “n.s.”.

## Results

### Astrocytic contributions to cortical synaptogenesis are differentially regulated between sexes

To elucidate sex differences in astrocyte-mediated synaptogenesis, we isolated cortical neurons and astrocytes with >95% purity from P1 Sprague-Dawley rat male or female pups. Neurons were cultured for 2 weeks in complete neuronal growth media (GM), with astrocytic factors provided by either astrocyte culture inserts (astro) or astrocyte-conditioned media (ACM) (Fig. 1A). Following immunocytochemical (ICC) staining, presynaptic (Bassoon) and postsynaptic (Homer1) puncta were imaged and their co-localization quantified to determine excitatory synapse number in the various treatment conditions. Both male and female-derived cortical neurons showed a significant increase in excitatory synapse density along their processes when cultured with sex-matched astrocyte inserts (Fig. 1B). The percent change (compared to GM only) of the astrocyte-induced synapses in female cultures was not as high as that in male cultures (221 ± 20% compared to 279 ± 28%, respectively), though this difference was not significant. To determine whether secreted factors from astrocytes differed in their synaptogenic potential depending on the sex of the donors, we treated male and female-derived cortical neurons with ACM from either male (M-ACM) or female (F-ACM) astrocytes. Excitatory synapse density was increased in neurons cultured from both sexes, and the magnitude of the increase was not different between M-ACM or F-ACM treatments (Fig. 1C). Interestingly, however, the percent change of the increase in synapses was significantly higher in male compared to female neurons for both M-ACM (207 ± 14% in male cultures compared to 157 ± 19% in female cultures) and F-ACM (218 ± 19% male, 162 ± 14% female; p<0.05, two-way ANOVA with Holm-Sidak’s *post hoc* analysis, *F*(_1,715_)=10.08). These results indicate that cortical neurons respond to astrocyte-secreted synaptogenic factors at different rates depending on their sex.

**Figure 1:**
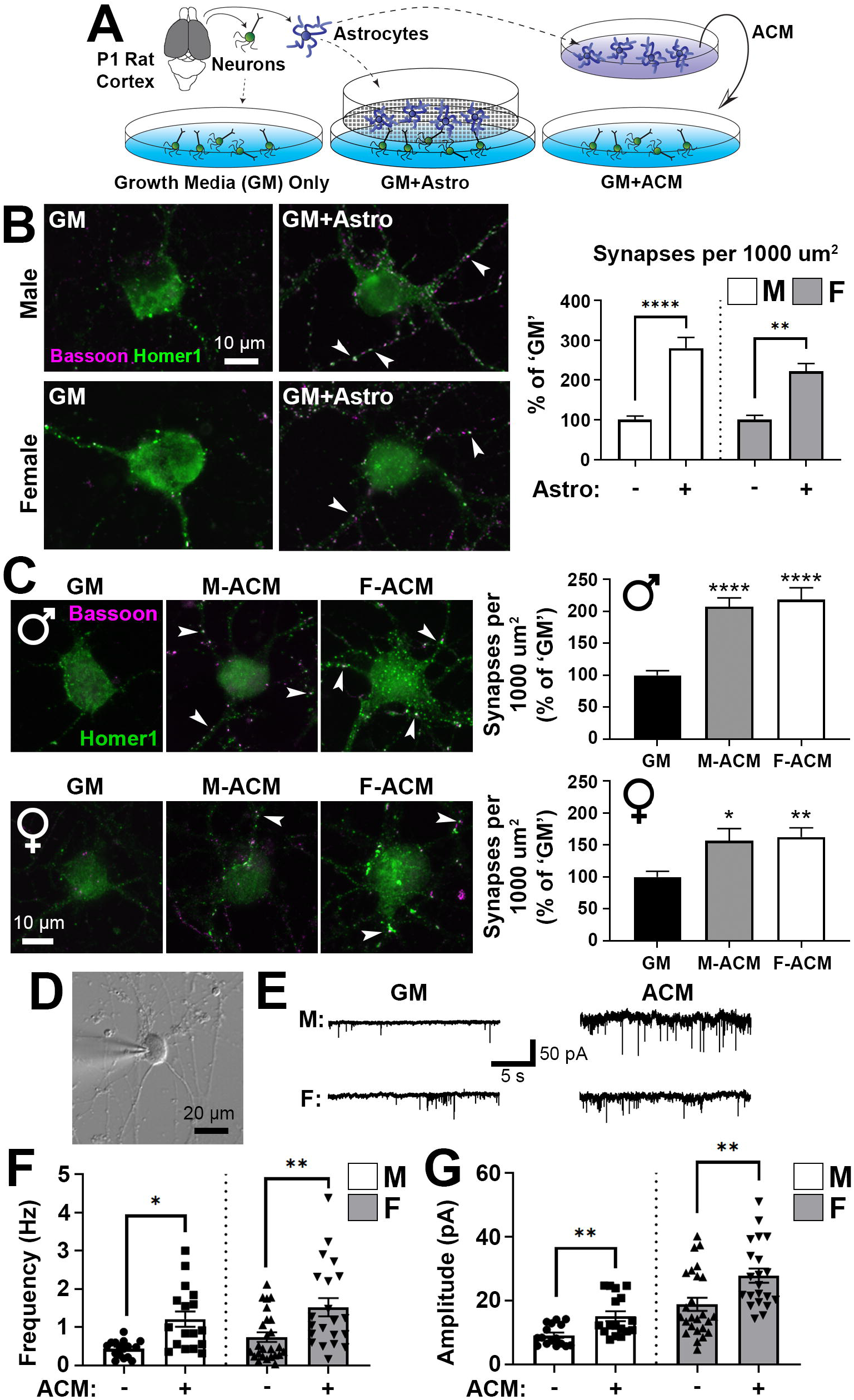
Sex differences in astrocyte-mediated synaptogenesis. **A,** Experimental model to study the contributions of astrocytes to synapse formation *in vitro*. Cortical neurons and astrocytes are purified separately from P1 male and female Sprague-Dawley rat littermates. Control conditions consist solely of neurons grown in complete neuronal growth media (“GM”). To test the impact of astrocyte-derived factors on synaptic development, neurons were treated with either astrocytes in cell culture inserts (“astro”) or astrocyte-conditioned media (“ACM”) prior to fixation and assessment via ICC. **B,** Left, Representative ICC images of cortical neurons isolated from either male or female pups and treated with either GM only or astrocyte inserts. Co-localized pre- (Bassoon; magenta) and postsynaptic (Homer1; green) puncta reveal sites of excitatory synapses (white arrowheads). Right, Excitatory synapse density expressed as a percentage of the sex-matched “GM” condition. **p<0.01, ****p<0.0001 (two-way ANOVA with Holm-Sidak’s *post hoc* analysis, *F*(_1,357_)=25.83; n=90 cells per condition from 3 independent experimental replicates). **C,** Left, Representative ICC images of male (top row) and female (bottom row)-derived cortical neurons treated with male or female-derived ACM (M-ACM and F-ACM, respectively), stained as in **B**. Right, Excitatory synapse density in male (top) and female (bottom)-derived neuronal cultures expressed as a percentage of the sex-matched “GM” condition. *p<0.05, **p<0.01, ****p<0.0001 (two-way ANOVA with Holm-Sidak’s *post hoc* analysis, *F*(_2,715_)=24.42; n=90-150 cells per condition from 3-5 independent experimental replicates). **D**, Brightfield image of a cortical neuron in whole cell patch clamp configuration for the recording of mEPSCs. **E,** Sample mEPSC traces from male (top) and female-derived (bottom) cortical neurons treated with (right) or without (left; “GM”) sex-matched ACM. **F** and **G,** Frequency (**F**) and amplitude (**G**) of mEPSCs recorded from male and female-derived cortical neurons treated with or without sex-matched ACM. *p<0.05, **p<0.01 (two-way ANOVA with Holm-Sidak’s *post hoc* analysis, *F*(_1,75_)=19.33 (**F**), *F*(_1,75_)=14.62 (**G**); n=16-25 cells per condition from 3 independent experimental replicates).

We next wanted to test whether astrocyte-induced synaptic activity was differently impacted by sex. Male and female cortical neurons were patched in whole-cell configuration and miniature excitatory postsynaptic currents (mEPSCs) were recorded to assess baseline synaptic activity (Fig. 1D-E). Both mEPSC frequency (Fig. 1F) and amplitude (Fig. 1G) increased in response to sex-matched ACM treatment in males and females. Comparison of fold-changes indicates a larger magnitude increase in frequency in male neurons compared to female (2.74 ± 0.46 compared to 2.01 ± 0.25, respectively), though this did not result in a significant difference. The fold-changes in amplitude increase were similar (1.61 ± 0.15 male, 1.46 ± 0.12 female), suggesting that potential sex differences in astrocyte-mediated excitatory synaptogenesis manifest primarily as changes in synapse number rather than strength.

### Thrombospondin promotes synaptic development in male but not female cortical neurons

To rule out the possibility that inherent sex differences in synapse density are responsible for the distinct effects of astrocyte-secreted factors on synaptic development, we compared basal synapse number between male and female-derived cortical neuron cultures via ICC (Fig. 2A). No differences were found in synaptic density between the sexes, indicating that the differences we previously observed must have been due to a differential response to astrocytic factors.

**Figure 2:**
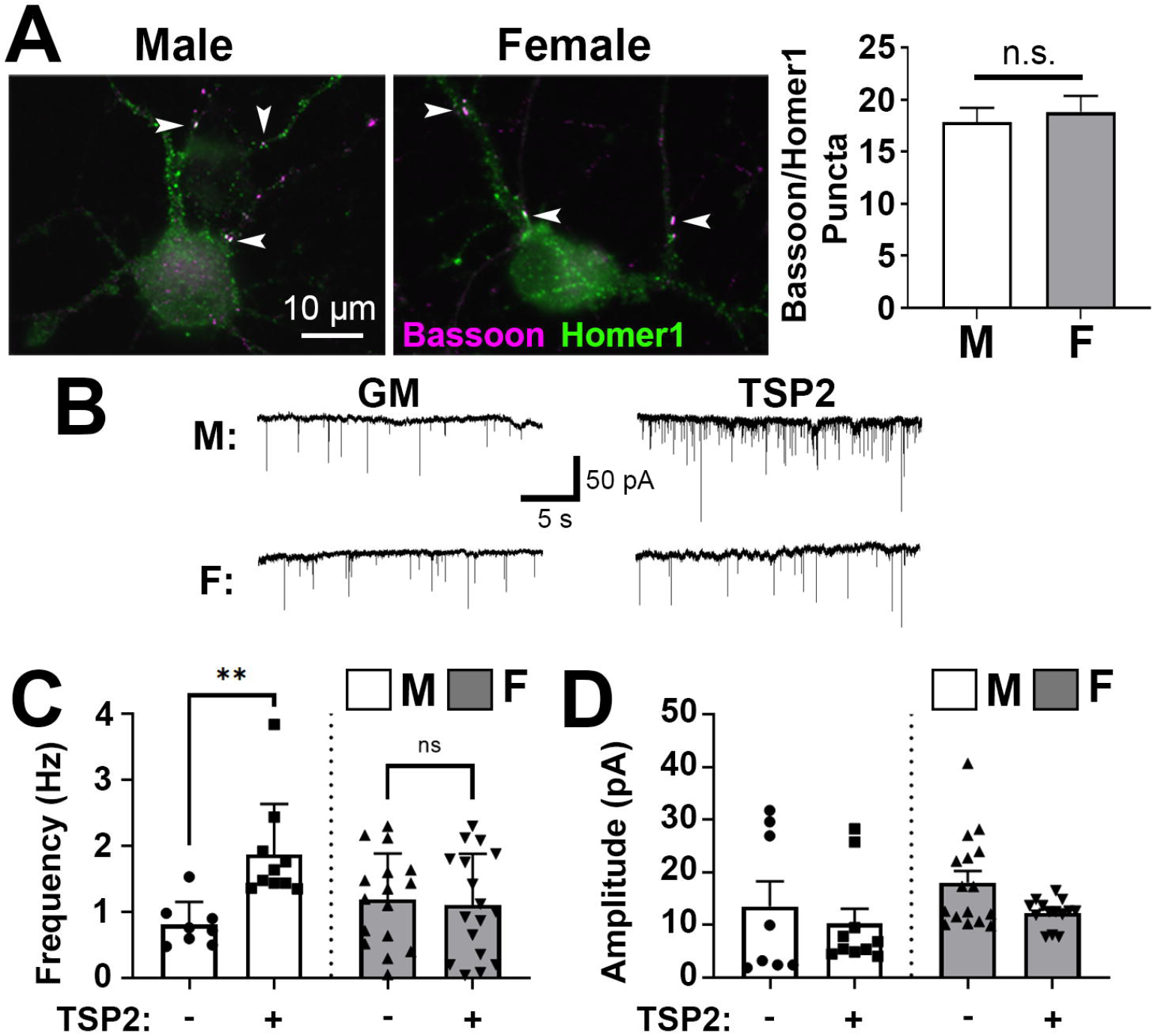
Increased synaptic response to astrocytic factor TSP by male but not female-derived cortical neurons. **A,** Left, Representative ICC images of male and female-derived cortical neurons. Co-localized pre- (Bassoon; magenta) and postsynaptic (Homer1; green) puncta reveal sites of excitatory synapses (white arrowheads). Right, Number of co-localized excitatory synaptic puncta. n.s.=not significant (two-tailed unpaired t-test; n=330 cells per condition from 11 independent experimental replicates). **B,** Sample mEPSC traces from male (top) and female-derived (bottom) cortical neurons treated with (right) or without (left; “GM”) purified TSP2. **C** and **D**, Frequency (**C**) and amplitude (**D**) of mEPSCs recorded from male and female-derived cortical neurons treated with or without TSP2. **p<0.01 (two-way ANOVA with Holm-Sidak’s *post hoc* analysis, *F*(_1,47_)=5.56 (**C**), *F*(_1,44_)=3.13 (**D**); n=8-17 cells per condition from 3 independent experimental replicates).

Thrombospondins (TSPs) were amongst the first astrocyte-secreted factors identified as having synaptogenic properties (Christopherson et al., 2005; Risher and Eroglu, 2012). However, this and other previous studies investigating the synapse-promoting properties of TSP were performed using neurons isolated from mixed-sex litters. In Risher et al. (2018), we conducted our *in vitro* studies in cortical neurons isolated specifically from male pups after determining that experimental success was highly variable when cultures were made from mixed-sex litters. To specifically test whether TSP has a synaptogenic effect in male but not female-derived cortical neurons, we next analyzed mEPSCs from male versus female cultures treated with purified TSP2 (Fig. 2B), one of the five synaptogenic TSP isoforms expressed in mammals. Intriguingly, only male-derived neurons showed increased mEPSC frequency after TSP2 treatment (Fig. 2C). Conversely, neurons in female cultures did not respond to TSP2 treatment with any changes in mEPSC frequency. Neither male nor female neurons showed any change in mEPSC amplitude following TSP2 treatment (Fig. 2D), but this finding was consistent with previous studies showing that TSPs promote the formation of NMDA-containing silent synapses that lack functional AMPA receptors (Risher and Eroglu, 2012). Nevertheless, these results indicate that TSP is synaptogenic in male but not female cortical neurons, providing a potential mechanism underlying the differential rates of astrocyte-induced synaptic development between the sexes.

### Modulation of TSP-induced synaptogenesis by estrogen

Why is TSP effective in promoting synapse formation in male but not female neurons? To begin to address this question, we next asked whether the unique responses of these cultures were due to differences in the sex hormone estrogen since, like astrocytes, sex hormones have been found to be powerful regulators of synaptic connectivity. Fluctuating levels of circulating estrogen, and androgens to a lesser extent, can drive significant changes in synaptic density in both sexes (Leranth et al., 2003; Srivastava and Penzes, 2011; Woolley and McEwen, 1992). Neurons, including those in dissociated culture, are capable of *de novo* production and secretion of estrogen (Hojo et al., 2004; Prange-Kiel et al., 2003). Konkle and McCarthy (2011) previously showed that cortical levels of 17β-estradiol (E2), the predominant biologically active form of estrogen, are transiently higher in female rats than males by the end of the first postnatal week. We observed a similar effect in our cortical neuron cultures, reaching significance at 9 days *in vitro* (DIV9; Fig. 3A). Interestingly, this time point corresponded with the peak period of synaptogenesis induced by TSPs, as well as the normal developmental expression of TSP1 and 2 (Christopherson et al., 2005; Risher and Eroglu, 2012). By contrast, astrocytes, which have also been identified as sources of estrogen in the brain (Azcoitia et al., 2011), did not show any sex differences in E2 secretion over this same time period (Fig. 3B). These findings suggested the possibility that neuronally-derived E2 may be a critical regulator of TSP-induced synaptogenesis.

**Figure 3:**
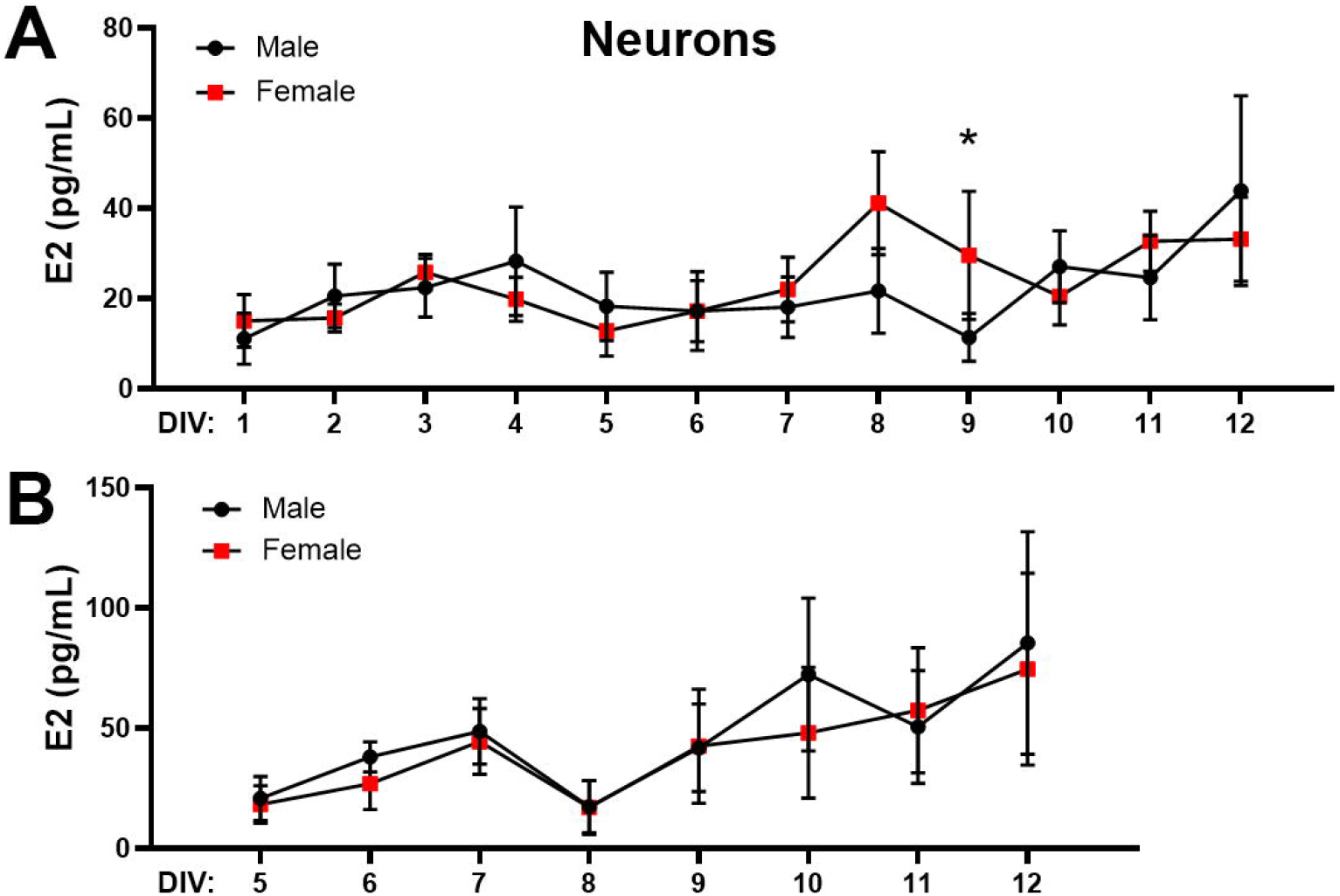
Transient increase in neuronally-derived estradiol in cultured female cortical neurons. **A,** Immunoassay results for levels of E2 detected in male versus female neuron-conditioned media. *p<0.05 (linear mixed-effects model [REML] with Holm-Sidak’s *post hoc* analysis, sex: *F*(_1,55_)=7.00; n=4 independent experimental replicates). **B,** Estradiol measurements from male versus female cortical ACM. n.s.=not significant (linear mixed-effects model, sex: *F*(_1,29_)=3.80; n=3 independent experimental replicates).

Estrogen is enzymatically converted from the hormone testosterone by aromatase. To investigate whether neuronally-synthesized estrogen interferes with TSP-induced synapse formation, we used ICC to quantify excitatory synapses in our purified cortical neuron cultures (Fig. 4A-B) following treatment with TSP2 and letrozole, an aromatase inhibitor. Following nearly 2 weeks of *in vitro* culture, ICC strengthened our electrophysiological findings (Fig. 2C) by showing that TSP2 significantly increased synapse numbers in male but not female-derived neurons (Fig. 4C). Intriguingly, treatment with letrozole revealed a sex-divergent effect of estrogen inhibition on the synapse-promoting ability of TSP2. While letrozole did not alter synapse density on its own, blockade of endogenous estrogen production did facilitate TSP2-induced synaptogenesis in female-derived neurons, albeit not to the same degree as in neurons isolated from males (Fig. 4C). The most unexpected result was seen in the male cultures, where letrozole seemed to have the opposite effect, greatly decreasing the magnitude of synaptogenesis induced by TSP2 (Fig. 4C). However, the combined effect of TSP2 and letrozole still led to increased synapses over the control treatment in male-derived cells, but now the percent change of this increase was virtually indistinguishable from that seen in the female cultures (164 ± 15% male, 160 ± 21% female). Taken together, these results demonstrate clear sex differences in TSP-induced synaptogenesis that may be regulated, at least in part, by estrogen synthesized endogenously by neurons.

**Figure 4:**
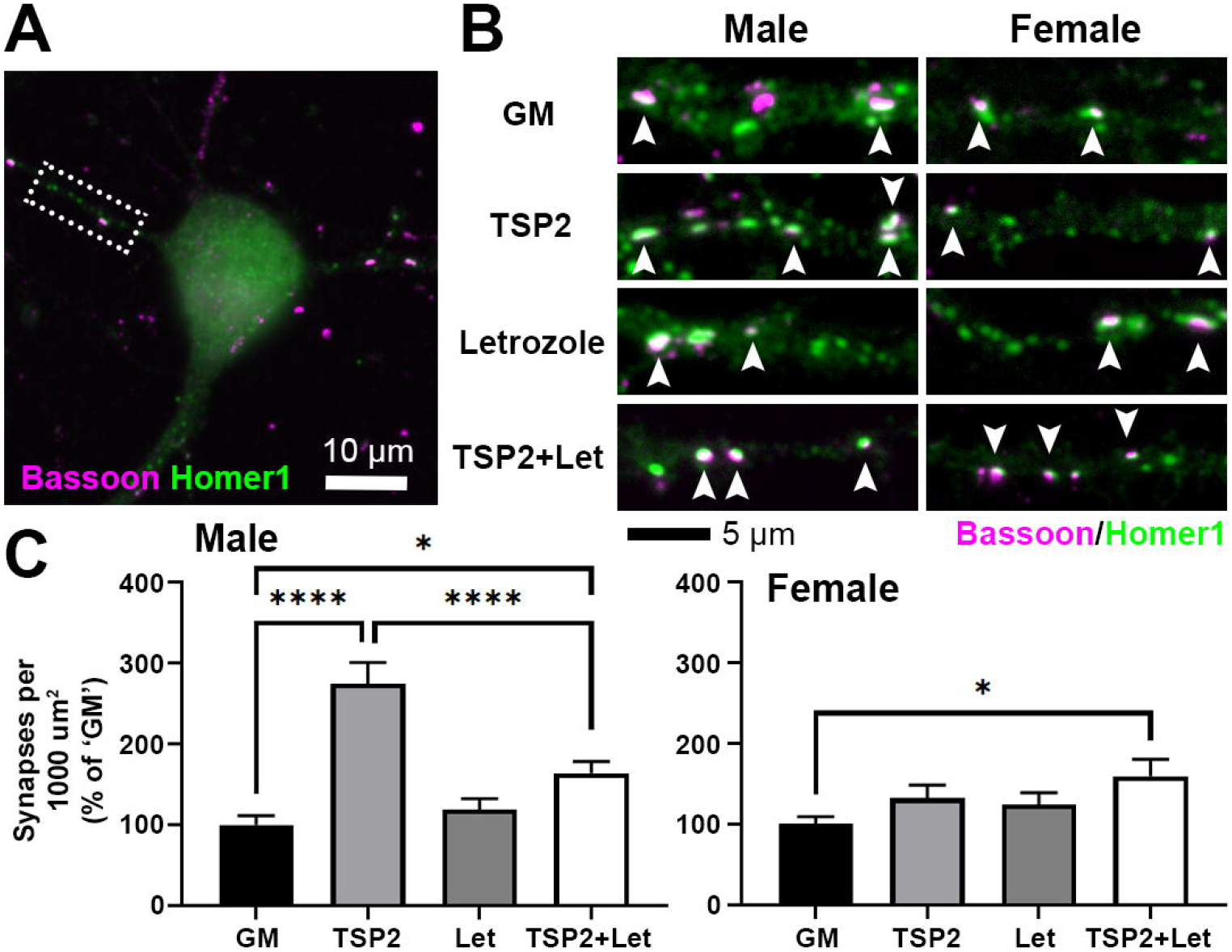
Synaptogenic properties of TSP are regulated by endogenous neuronal estrogen. **A,** ICC image of a male-derived cortical neuron stained with pre- (Bassoon; magenta) and postsynaptic (Homer1; green) markers. Dotted box indicates example dendritic region of interest sampled for images in **B,** which shows co-localized excitatory synaptic puncta (white, arrowheads) along male and female-derived cortical dendrites treated with GM only or GM plus TSP2, letrozole (Let), or TSP2/letrozole. **C,** Excitatory synapse density in male (left) and female (right) neuronal cultures expressed as a percentage of the sex-matched “GM” condition. *p<0.05, ****p<0.0001 (two-way ANOVA with Holm-Sidak’s *post hoc* analysis, *F*(_3,804_)=18.05; n=60-150 cells per condition from 2-5 independent experimental replicates).

## Discussion

Female and male brains show fundamental differences throughout development and evolution, with the adult human female brain estimated to contain approximately two-thirds the number of synapses across all cortical layers compared to males (Alonso-Nanclares et al., 2008). Though synapse number should certainly not be considered an indicator of intelligence or predictive of competency at any particular task, this difference does raise the possibility of fundamentally sex-distinct mechanisms of how synaptic networks are formed, developed, and maintained. Here, we provide evidence for one such mechanism, whereby cortical neurons purified from males, but not females, are highly receptive to the astrocyte-secreted synaptogenic factor TSP. Disruption of neuronal estradiol synthesis abolished this sex difference and resulted in TSP being weakly synaptogenic for both sexes, highlighting endogenous brain estrogen as a potentially critical player in astrocyte/synaptic signaling.

Thrombospondin is a potent synaptic organizer secreted by astrocytes to act through the neuronal L-type calcium channel auxiliary subunit, α2δ-1, during a peak period of synaptogenesis (Eroglu et al., 2009; Risher and Eroglu, 2012). This period consists of roughly the second to third postnatal weeks of rodent brain development, coinciding with a stage of rapid astrocyte growth and elaboration (Farhy-Tselnicker and Allen, 2018) that is thought to be roughly correlated with perinatal human brain development (Semple et al., 2013). Since this time point falls well before the onset of puberty (typically between days 34-48 in rats), neurosteroids have not traditionally been considered to play a significant role in shaping synaptic connectivity. Though we cannot rule out potential contributions from the gonadal testosterone surge near the end of gestation in male rats (Weisz and Ward, 1980), our observation of significant sex differences prior to the developmental milestone of gonadal hormone release, and the elimination of this difference by letrozole, suggests that our findings are likely due to *de novo* estrogen synthesis occurring in the very young brain. The phenomenon of *de novo* estrogen production by brain cells has previously been reported (McCarthy, 2008), but the importance of this process for proper neural circuit formation and operation had not been determined. Our results so far certainly indicate that neuronally-sourced estradiol regulates astrocyte-mediated synaptic development, though defining the precise nature of this role has been complicated by the seemingly contradictory finding that blocking aromatase reduces the synaptic capability of TSP in male-derived neurons yet is permissive in female cultures. Studies are ongoing to elucidate this mechanism more fully, as well as to establish the relevance of this process for cortical development *in vivo*.

The role of estrogen as a master regulator of synaptic connectivity has been well-established (see Grkovic and Mitrovic (2020) for review). However, many previous studies in the field that established the effects of estrogen on synapse formation were performed in conditions that included glia, such as intact brain, hippocampal slice culture, or dissociated neuronal culture with up to 20% glial contamination (Brandt and Rune, 2019; Kramar et al., 2009; Kretz et al., 2004; Lu et al., 2019; Mukai et al., 2007; Zhao et al., 2018). Both neurons and astrocytes express estrogen receptors (Hosli and Hosli, 1999; Hosli et al., 2001), while astrocytes in particular can rapidly respond to fluctuating levels of estrogen (Chaban et al., 2004), suggesting a potential confound for these findings. In our neuronal cultures, we achieve 95% purity (confirmed by ICC) and include cytosine arabinoside (AraC) to inhibit the proliferation of the astrocytes that are present. In this way, we are able to isolate the responses of neurons and specifically provide the factors typically sourced from astrocytes, either as individual components (i.e. TSPs) or as a heterogeneous mixture (i.e. ACM). As the findings in this study have indicated, such an approach may be necessary to continue unraveling the intersection between estrogen and astrocytes in the shaping of synaptic connectivity (McCarthy et al., 2003).

In addition to the known differences between male and female brains of healthy individuals, aberrant brain states are also known to manifest differently between the sexes. Diseases of the aging brain, such as Alzheimer’s and Parkinson’s, present with highly different rates of incidence and severity between men and women, while many more boys than girls are diagnosed with certain neurodevelopmental disorders (NDDs) including schizophrenia and autism spectrum disorder (Abel et al., 2010; Hanamsagar and Bilbo, 2016; Werling and Geschwind, 2013). In the case of NDDs, it is tempting to speculate that early dysregulation of synaptic connectivity may contribute to disease pathogenesis in a sex-dependent manner. Indeed, synaptic pathology and aberrant spinogenesis are common findings in both autism and schizophrenia, with a number of overlapping candidate molecules between the disorders (including TSP receptor α2δ-1 and its associated calcium channel subunits) that suggest some degree of common mechanistic dysfunction (Harrison et al., 2019; Iossifov et al., 2014; Penzes et al., 2011; Risher et al., 2018). Future studies that aim to elucidate disease pathology for the sake of developing novel, targeted therapies (Gillies and McArthur, 2010) should therefore take into consideration whether astrocyte-mediated synaptic signaling pathways are differentially regulated and/or disturbed in one sex compared to the other.

In summary, our findings have revealed that at least one prominent synaptogenic pathway regulated by astrocytes displays a significant sex bias, with thrombospondin-2 promoting synapse formation in male but not female cortical neurons. Future studies investigating other astrocyte factors, such as TSPs 1 & 4, hevin (Risher et al., 2014), SPARC (Kucukdereli et al., 2011), glypicans (Allen et al., 2012; Farhy-Tselnicker et al., 2017), and TGF-β (Diniz et al., 2014), among others, may determine whether this phenomenon affects other glial signaling pathways or is specific to TSP. From this starting point, we can start to more thoroughly investigate how astrocytes promote synaptic connectivity in both sexes, elucidating potential mechanisms that underlie fundamental differences in the male and female brain to address an understudied yet vitally important area of neuroscience (McCarthy et al., 2012).

## Acknowledgements

We would like to thank Dr. Kelly Hopper and the staff of the Marshall University Animal Resource Facility for excellent veterinary assistance.

This research was supported by MU startup funds to B.J.H. and W.C.R., the MU Undergraduate Creative Discovery & Research Scholar Award to E.H.B., the Marshall University Genomics and Bioinformatics Core, the West Virginia IDeA Network of Biomedical Research Excellence (WV-INBRE) grant (P20GM103434), the COBRE ACCORD grant (1P20GM121299), and the West Virginia Clinical and Translational Science Institute (WV-CTSI) grant (2U54GM104942).

Author contributions: W.C.R. designed the study. A.M., E.H.B., B.J.H., and W.C.R. conducted the experiments. A.M., B.J.H., and W.C.R. analyzed the data. A.M. and W.C.R. wrote the manuscript.

